# Metagenomic analysis of virus diversity and relative abundance in a eutrophic freshwater harbour

**DOI:** 10.1101/690891

**Authors:** Christine N. Palermo, Roberta R. Fulthorpe, Rosemary Saati, Steven M. Short

## Abstract

Aquatic viruses have been extensively studied over the past decade, yet fundamental aspects of freshwater virus communities remain poorly described. Our goal was to characterize particle-associated virus communities seasonally and spatially in a freshwater harbour. Community DNA was extracted from water samples and sequenced on an Illumina HiSeq platform. Assembled contigs were annotated as belonging to the virus families *Caudovirales, Mimiviridae, Phycodnaviridae*, and virophages (*Lavidaviridae*), or to other groups of undefined viruses. Diverse *Mimiviridae* contigs were detected in the samples, but the two sites contained distinct *Mimiviridae* communities. Virophages were often the most abundant group, and discrete virophage taxa were remarkably stable across sites and dates despite fluctuations in *Mimiviridae* community composition. *Caudovirales* were present at low abundances in most samples, contrasting other studies of freshwater environments. Similarly, *Phycodnaviridae* abundances were surprisingly low in all samples despite the harbour’s capacity to support high algal biomass during the summer and autumn months, suggesting that *Mimiviridae* are the dominant algae-infecting viruses in this system. Overall, our findings provided insights into freshwater virus community assemblages by expanding the documented diversity of freshwater virus communities, highlighting the potential ecological importance of virophages, and revealing distinct communities over small spatial scales.

## INTRODUCTION

Viruses can modify and control the structure and function of ecosystems and in turn, influence global biogeochemical cycles and the evolution of organisms [1]. Historically, use of traditional culture-based methods to study environmental viruses led to underestimations of their abundance and ecological importance [2], whereas more contemporary molecular methods allowed analyses of environmental microbial communities without the constraints of cultivation. However, unlike the prokaryotes and eukaryotes they infect, viruses do not share universally conserved genes that can be readily targeted to survey entire communities. Thus, viral ecology is a field for which major advances can be realized through shotgun metagenomic sequencing [3]. Nonetheless, despite intensive efforts over the past couple of decades, comprehensive knowledge of virus community diversity and dynamics remains elusive for most natural settings.

In the first viral metagenomics study, over 65% of sequences recovered from surface seawater samples were not significantly similar to any sequence in existing databases, highlighting the lack of knowledge of environmental viruses [4]. Since then, viral sequence databases have expanded dramatically, in large part due to several large-scale ocean sampling expeditions that have included viral community analysis (as reviewed in [5]). These sampling expeditions include the Tara Oceans Expedition, the Malaspina Circumnavigation Expedition, Pacific Ocean Virome (POV), the San Pedro Ocean Time-series (SPOT), and the Bermuda Atlantic Time-series Study (BATS). BATS tracked viral abundance in the Sargasso Sea over a decade, while the SPOT studies measured temporal variation in virus communities and their hosts. The POV was established with data from transects spanning from coastal waters to the open ocean to document spatial changes in microbial and virus communities, whereas the Tara Oceans and Malaspina Circumnavigation Expeditions were designed to gather baseline global oceanic biodiversity data.

The Tara Oceans Expedition was conducted from 2009-2013 with the aim of globally sampling a wide range of organismal and functional diversity in the surface oceans, while the Malaspina Circumnavigation Expedition sailed from December 2010 to July 2011 with a focus on deep ocean microbiology. The Tara Oceans Expedition resulted in several important discoveries related to diverse groups of marine planktonic taxa, including viruses (e.g. [6-10]). Importantly, this sampling expedition provided data supporting previous observations of high local diversity but limited global diversity that led to the conception of a ‘seed-bank’ model of virus diversity [11]. The Tara Oceans survey revealed that virus community composition was strongly impacted by temperature and oxygen concentrations on local scales due to these factors’ influence on their hosts, while on larger scales ocean currents were responsible for transporting and mixing a virus ‘seed-bank’ [6]. Furthermore, using 17 viromes generated from the expedition, the abundance and diversity of nucleo-cytoplastic large DNA viruses (NCLDVs) were mapped revealing that there were approximately 10^4^-10^5^ viruses per ml in the photic zone and that the so-called ‘Megavirales’ and the *Phycodnaviridae* were the most common NCLDVs in the epipelagic oceans [12]. Complementing the Tara Oceans Expedition, data stemming from the Malaspina Circumnavigation Expedition demonstrated that viruses have higher turnover rates in the deep ocean compared surface waters, and they play important roles in DOC production and nutrient release, especially in the bathypelagic [13]. By combining viral sequences from the Tara Oceans and the Malaspina Circumnavigation Expeditions, numerous virus genomes have been assembled, expanding viral sequence databases more than three-fold [14]. Thus, these expeditions have vastly improved our understanding of marine viral ecology and have highlighted the global importance of virus activity.

Though extensive surveys of marine virus communities have been conducted, relatively little is known about fundamental aspects of freshwater virus ecology, such as their distribution in the environment. Despite their underrepresentation in databases, research has demonstrated that freshwater virus communities contain novel viruses and are distinct from other aquatic virus communities [15-17]. Metagenomics has been used to study virus communities in natural freshwater lakes from the Arctic [18], Canada [19], USA [20,21], Ireland [22], France [17], China [23], and Antarctica [24,25]. With respect to virus communities in eutrophic lakes, studies by Green et al. [20], Skvortsov et al. [22], and Ge et al. [23] revealed virus communities dominated by *Caudovirales* in the epilimnion of eutrophic lakes in China, USA, and Ireland, respectively, but other dsDNA viruses, unclassified bacteriophages, and ssDNA viruses were also detected albeit at lower abundances. Roux et al. [26] focused on the virophage and NCLDV communities in a eutrophic freshwater lake in USA, and observed highly dynamic virophage communities lacking any apparent annual or seasonal patterns of abundance. Although viral sequence databases have expanded rapidly alongside knowledge of marine virus ecology, documenting virus diversity and activity in disparate freshwater environments remains an exciting and untapped area for research.

The goal of the research described here was to characterize the virus community in Hamilton Harbour, a eutrophic freshwater embayment of Lake Ontario. The harbour is located at the western end of Lake Ontario and has an area of 21.5-km^2^ and an average depth of 13 m. It is separated from Lake Ontario by a naturally occurring sandbar and the Burlington Shipping Canal [27], and is the largest Canadian port in the Great Lakes [28]. The area surrounding the harbour has a long history of industrial activity, resulting in a highly polluted harbour containing heavy metals and many other hazardous contaminants [29,30]. Moreover, inputs from wastewater treatment plants, stormwater and sewage overflow, and agricultural and urban runoff have led to high nutrient concentrations, especially phosphorus. Thus, Hamilton Harbour is a eutrophic system that experiences seasonal blooms of cyanobacteria and algae, poor water clarity, and depleted hypolimnetic oxygen. As a result of all these perturbations, the harbour was designated an ‘Area of Concern’ in the amended 1987 USA-Canada Great Lakes Water Quality Agreement (GLWQA). Despite its designation over 30 years ago and ongoing remediation efforts, Hamilton Harbour remains one of the most impaired sites in the Canadian Great Lakes [31].

While Hamilton Harbour is in general an extensively studied system, the microbial community has only been examined using microscopic techniques to investigate microbial diversity and abundance. To our knowledge, there are no published studies characterizing the microbial community in Hamilton Harbour using metagenomics, nor are there any studies of the virus community in Hamilton Harbour. The metagenomic dataset used in this study was originally generated to study Hamilton Harbour bacteria and eukaryotic plankton, but was leveraged herein to provide an initial assessment of the cell- and particle-associated virus community and to determine if seasonal and spatial patterns of diversity and abundance could be discerned in this unique freshwater environment. More research is required to better characterize freshwater virus diversity, community structures, and patterns of abundance, and the factors that drive these phenomena. This research aims to address these knowledge gaps by providing a detailed view of the virus community in Hamilton Harbour.

## MATERIALS AND METHODS

### Sampling Sites and Collection

Water samples were collected from two long-term Environment Canada monitoring sites in Hamilton Harbour (referred to as stations 1001 and 9031 in previous publications). One site (station 1001) is located at the deepest and most central part of the harbour (43°17’17.0”N 79°50’23.0”W) with a water depth of 24 m and a 1.2 km distance from the shoreline. The other site (station 9031) is located less than 0.5 km from the shoreline (43°16’50.0”N 79°52’32.0”W), with a water depth of 12 m. This “nearshore” site is influenced by effluents from the Cootes Paradise watershed on the west end of the harbour, while the “mid-harbour” site is closer to the Burlington Shipping Canal and Lake Ontario. In 2015, water samples for metagenomic analyses were collected on July 30^th^, August 13^th^ and 27^th^, and September 10^th^ and 24^th^, from both sites at approximately 1 m below the surface using a Van Dorn bottle sampler; in total, 10 samples were collected, 5 from each of the nearshore and mid-harbour sites. 500 ml water samples were filtered through 0.22 μm pore-size Sterivex capsule filters (EMD Millipore, USA), and the filters were sealed and stored at −80°C until further analysis. For each sample collected, physiochemical parameters including pH, dissolved oxygen, temperature, redox potential, and chlorophyll a were measured *in situ* using a YSI 58 (Xylem Inc., USA). Secchi depth was also measured with each sample.

### DNA Extraction and Sequencing

To extract community DNA from each sample, biomass was recovered from the filters by adding 2 ml of molecular grade nuclease free water to each filter and vortexing for 5 minutes. The resuspended material was transferred to a 1.5 ml microcentrifuge tube under sterile conditions and was centrifuged at 6,000 x g for a total of 15 minutes. DNA was extracted from the pelleted material using a FastDNA SPIN Kit (MP BIO, USA), beginning with the addition of 500 μl of CLS-Y solution. The manufacturer’s protocol was followed except the wash step was repeated three times to maximize removal of environmental contaminants. Following extraction, DNA concentrations were estimated using a NanoDrop ND-1000 UV-Vis Spectrophotometer (NanoDrop Technologies, USA), and were standardized to 1.5 μg of DNA per sample before being sent for library preparation and shotgun metagenome sequencing by MR. DNA (Molecular Research LP, USA).

Sample DNA libraries were prepared using a Nextera DNA Sample Preparation Kit (Illumina, USA), following the manufacturer’s protocol. DNA concentrations were measured using the Qubit dsDNA HS Assay Kit (Life Technologies, USA) before and after library preparation (Table 1). Prior to library preparation, samples were diluted to achieve the recommended concentration of 2.5 ng μl^-1^. For the August 27 nearshore and mid-harbour sites, achieving a concentration of 2.5 ng μl^-1^ was not possible, so the maximum volume (20 μl) of each sample was used for these libraries. Average library size was determined using an Agilent 2100 Bioanalyzer (Agilent Technologies, USA). Each library was clustered using a cBot System (Illumina, USA), and was sequenced from paired ends using 500 cycles with the HiSeq 2500 (2 x 250 bp) system (Illumina, USA).

**Table 1:** Initial DNA concentration, final library concentration, and average fragment size in the library for each of the 10 samples in this study.

| Sample | Initial DNA Concentration (ng/μl) | Library Concentration (ng/μl) | Average Library Size (bp) |
| --- | --- | --- | --- |
| July 30 – Nearshore | 5.30 | 16.2 | 985 |
| July 30 – Mid-harbour | 2.84 | 18.3 | 1070 |
| August 13 – Nearshore | 2.76 | 17.5 | 1070 |
| August 13 – Mid-harbour | 3.40 | 14.5 | 1000 |
| August 27 – Nearshore | 2.28* | 15.0 | 950 |
| August 27 – Mid-harbour | 1.42* | 8.94 | 615 |
| September 10 – Nearshore | 9.58 | 12.0 | 1000 |
| September 10 – Mid-harbour | 10.6 | 14.1 | 1060 |
| September 24 – Nearshore | 9.10 | 15.7 | 1030 |
| September 24 – Mid-harbour | 2.68 | 17.7 | 900 |

\* denotes that samples could not be adjusted to 2.5 ng/μL prior to library preparation.

### Metagenome Data Processing

After sequencing and base-calling, adapters and barcodes were removed by MR DNA (Molecular Research LP, USA). Read quality parameters were verified using FastQC version 0.11.5 [32] prior to quality control and once again prior to assembly. Quality control was performed using a sliding window method with the program Sickle version 1.33 [33] using a quality score cut-off value of 30. All reads shorter than 50 bp were removed prior to assembly. Reads were assembled using IDBA-UD version 1.1.3 [34] with alterations to the source code to accommodate longer read lengths and higher maximum k values (as in [22]). The de Bruijn graph-based assembly was performed using k values from 20 to 200 in increments of 20. A minimum k-mer count of 1 was used in the assembly to maximize assembly of the low coverage reads. Contig alignment was achieved with BLASTx against the April 2018 NCBI-nr database downloaded from ftp://ftp.ncbi.nih.gov/blast/db/FASTA/nr.gz. DIAMOND version 0.9.19 [35] was used for alignment with frameshift alignment and very sensitive modes activated. MEGAN6-LR version 6.11.4 [36] was used to annotate contigs using the Lowest Common Ancestor (LCA) algorithm in long read mode with a bit score cut-off value of 100 and a 10^-6^ e-value cut-off. The March 2018 MEGAN protein accession mapping file was downloaded from http://ab.inf.uni-tuebingen.de/data/software/megan6/download/welcome.html. Quality controlled reads were mapped back to assembled contigs using Bowtie 2 [37] in very sensitive mode, and mapping information for each contig was extracted using SAMtools [38]. Table 2 summarizes the number of reads and contigs at each step in the pipeline for each sample.

**Table 2:** Summary of the number of reads and contigs at each step in the data processing pipeline

| Sample | Reads pre-QC | Reads post-QC | Contigs post-assembly (Mean length) | Contigs assigned | Virus contigs assigned | Percent of virus contigs |
| --- | --- | --- | --- | --- | --- | --- |
| July 30 – Nearshore | 13,403,832 | 11,608,892 | 480,233 (693) | 212,156 | 349 | 0.16 |
| July 30 – Mid-harbour | 13,206,316 | 11,246,470 | 433,927 (824) | 256,277 | 397 | 0.15 |
| August 13 – Nearshore | 12,758,364 | 11,040,814 | 398,774 (741) | 196,128 | 488 | 0.25 |
| August 13 – Mid-harbour | 13,104,992 | 11,414,576 | 416,468 (821) | 273,693 | 662 | 0.24 |
| August 27 – Nearshore | 15,685,750 | 13,953,600 | 374,464 (662) | 128,233 | 484 | 0.38 |
| August 27 –<br>Mid-harbour | 16,101,196 | 14,751,904 | 233,868<br>(834) | 183,817 | 232 | 0.13 |
| September 10 –<br>Nearshore | 16,228,148 | 14,257,158 | 418,490<br>(609) | 110,567 | 463 | 0.42 |
| September 10 –<br>Mid-harbour | 15,411,142 | 13,561,346 | 443,692<br>(568) | 64,142 | 604 | 0.94 |
| September 24 –<br>Nearshore | 14,689,040 | 12,764,018 | 367,778<br>(629) | 112,144 | 372 | 0.33 |
| September 24 –<br>Mid-harbour | 13,151,390 | 11,228,970 | 296,813<br>(777) | 185,913 | 104 | 0.06 |

### Metagenome Data Analysis

Once taxonomic assignments were applied to contigs, their relative abundances were estimated after normalizing by contig length and sequencing depth per sample. The number of reads that mapped to each contig was divided by the contig length in kbp. These values were summed for all contigs in a sample and were divided by a normalizing factor of 1000. Then, the number of mapped reads per kbp was divided by the normalized sum to achieve a relative abundance count that could be compared to other groups within and between samples. All virus contig assignments were sorted into one of the following categories based on the NCBI taxonomic classifications: *Caudovirales, Mimiviridae, Phycodnaviridae*, virophages (*Lavidaviridae*), *Iridoviridae, Poxviridae*, other dsDNA viruses, ssDNA viruses, unclassified bacterial viruses, and unclassified viruses. Because the assigned taxonomic classifications for some contigs in the “unclassified bacterial viruses” and “unclassified viruses” categories were less specific than information provided in the literature within which they were originally reported, some contigs in these categories were manually curated and were assigned to more specific groups based on published information (Table S1).

As well as assigning some contigs to more specific categories, some of the more specific assignments in the dsDNA viruses group were re-assigned to the “Other dsDNA viruses” group if they were observed in less than half of the samples and at <0.5% abundance. For example, contigs annotated as *Marseilleviridae* were only observed in 2 of the 10 samples at < 0.2 % abundance. Because there was not enough representation to making meaningful comparisons, the *Marseilleviridae* were grouped together with the “Other dsDNA viruses”. As a final note, in one sample a single contig was annotated as an RNA virus, and this contig was re-assigned to the “Unclassified viruses” category (Table S1).

There is evidence that some viruses previously and tentatively considered phycodnaviruses (i.e., members of the *Phycodnaviridae* family) are more closely related to *Mimiviridae* than *Phycodnaviridae*, and in fact form a separate subfamily ‘Mesomimivirinae’ within the *Mimiviridae* [39]. Though not formally recognized by the International Committee for Taxonomy of Viruses (ICTV), several publications have used the terms ‘*Megaviridae*’ or ‘extended *Mimiviridae*’ to refer to ‘Mesomimivirinae’ [40]. Here we will use the term *Mimiviridae* to encompass the formally recognized *Mimiviridae* as well as the proposed ‘extended *Mimiviridae*’ subfamily that are often considered phycodnaviruses. Proposed members of the ‘Mesomimivirinae’ include Organic Lake phycodnaviruses (OLPVs), *Aureococcus anophageefferens* virus (AaV), *Chrysochromulina ericina* virus (CeV), *Phaeocystis pouchetii* virus (PpV), *Pyramimonas orientalis* virus (PoV), and Group I *Phaeocystis globosa* viruses (PgVs) [39,41-44]. Based on these recent developments, we manually curated the affiliation of viruses that belong to the proposed Mesomimivirinae group but were annotated as *Phycodnaviridae* in the current NCBI scheme (accessed April 2018), as viruses within the *Mimiviridae* (Table S1).

A boxplot of the overall virus community was generated using the “ggplot2” package [45] in RStudio. Bray-Curtis dissimilarity with unweighted pair group method with arithmetic mean (UPGMA) clustering was used to determine which samples were most similar based on relative abundances of different virus groups. The relationship between the relative abundance of virus groups, environmental parameters (dissolved oxygen, pH, chlorophyll a, temperature, Secchi depth, and redox potential), and sites, was statistically tested using canonical correspondence analysis (CCA); tests of 10,000 permutations were used to compute the significance of the model and the variables. Cluster analysis and CCA were computed using the “vegan” package [46] in RStudio. For samples collected on July 30^th^, and September 10^th^ and 24^th^, data were available for other factors including: ammonia, chloride, fluoride, sulfate, particulate organic carbon, particulate organic nitrogen, nitrate and nitrite, total dissolved nitrogen, total phosphorus, total dissolved phosphorus, total particulate phosphorus, and soluble reactive phosphorus. Separate CCA models were tested for the entire dataset and the subset of dates for which additional data were available.

## RESULTS

### Virus community composition in Hamilton Harbour

Diverse virus contigs were identified in Hamilton Harbour and some were classified within the virus families *Caudovirales, Mimiviridae, Phycodnaviridae*, virophages (*Lavidaviridae*), while others could only be classified as belonging to *unclassified bacteriophages, other dsDNA viruses*, and *ssDNA viruses*. Henceforth, for the sake of brevity, we will name only virus groups when referring to contigs annotated as viruses from those groups. Overall, virophages were the most abundant (44.7% average abundance across all samples) but were also the most variable (from 0.1% to 74.5% abundance). The *Mimiviridae* were the second most abundant group, comprising an average of 21.1% of the virus community across all samples. Interestingly, though they are intimately associated, the abundances of *Mimiviridae* (ranging from 8.2% to 36.0% abundance) did not fluctuate to the same extent as the virophages (Figure 1). At the nearshore site, virophages represented a large percentage of the virus community in every sample with abundances between 47.4% and 71.7%, yet their abundances fluctuated widely at the mid-harbour site ranging from 0.12% to 74.5%. *Mimiviridae* were consistently detected as a substantial proportion of the community in the nearshore samples with relative abundances ranging from 18.2% to 34.6% overall. Again, like the virophages, *Mimiviridae* abundances were more variable at the mid-harbour site, comprising between 8.2% and 36.0% of all virus contigs. Similarly, *Caudovirales* consistently represented less than 5.0% of the virus community in the nearshore samples, but ranged from 1.3% to 73.9% at the mid-harbour site. *Phycodnaviridae* were a minor component of the virus community in all samples, representing only 0.09% to 3.3% at the nearshore site and 0.05% to 1.9% at the mid-harbour site (Figure 2).

**Figure 1:**
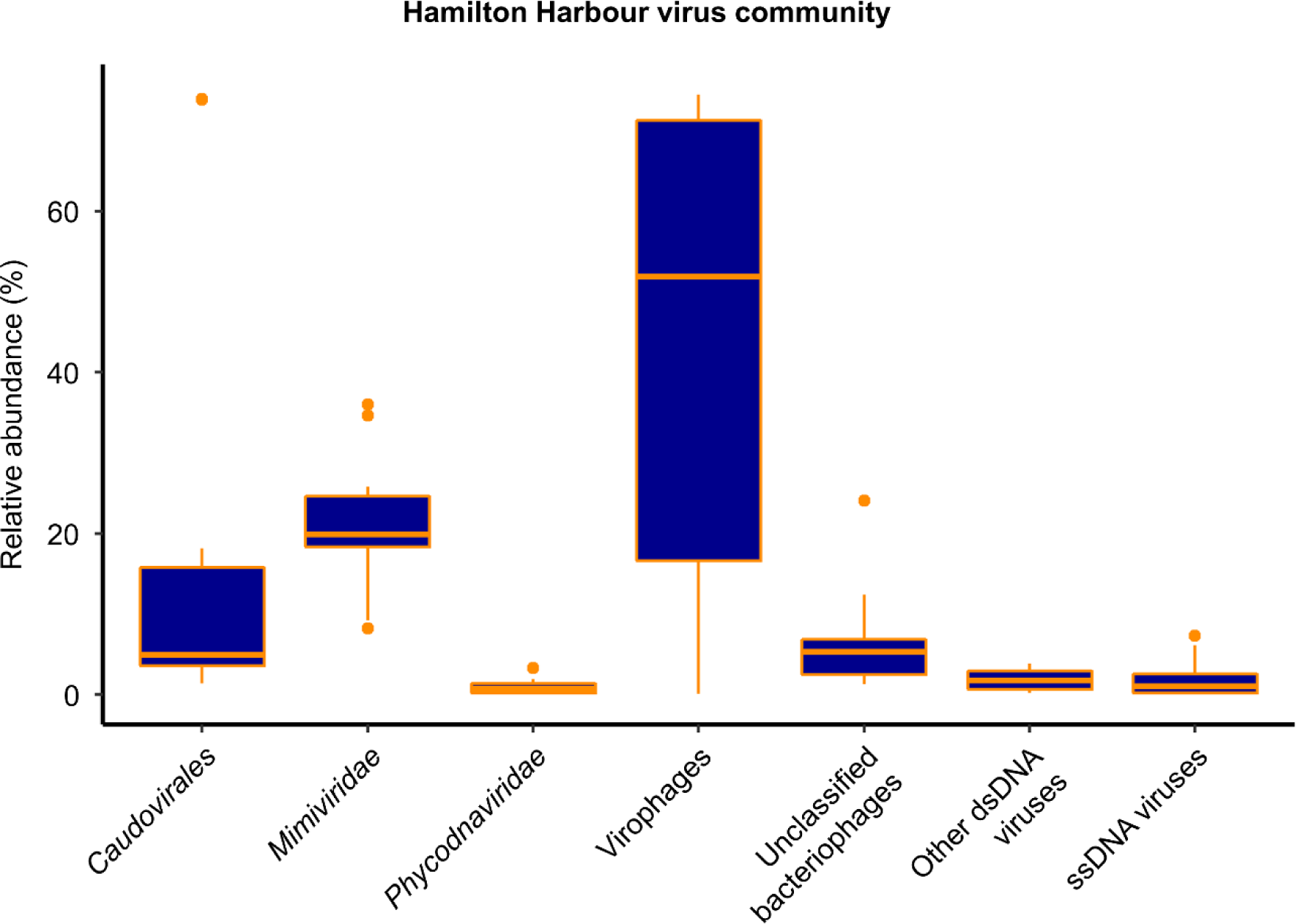
Boxplot of the overall virus community in Hamilton Harbour based on all 10 metagenomes from the 5 sampling dates at the nearshore and mid-harbour sites.

**Figure 2:**
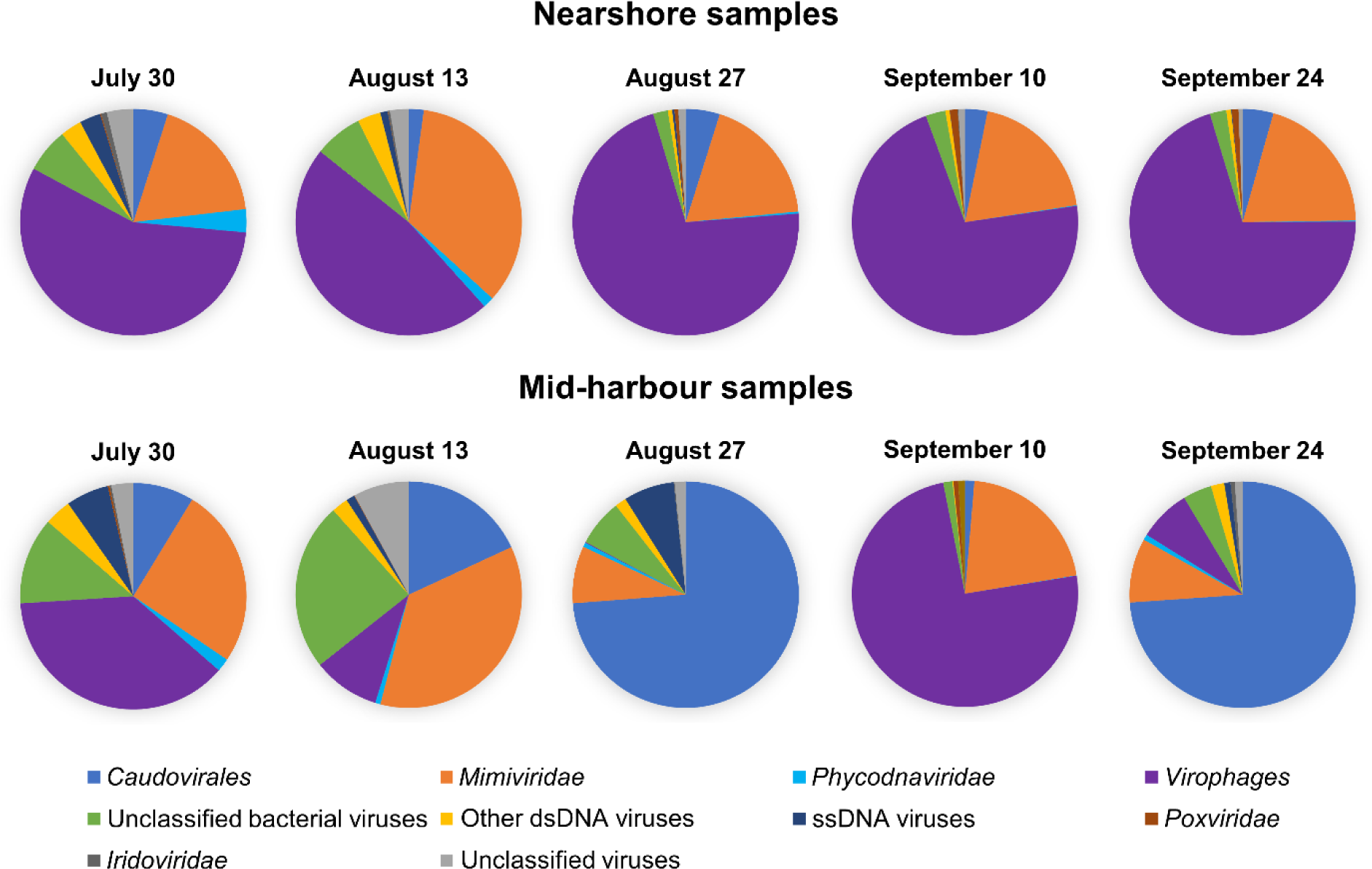
Individual virus communities for each of the 10 metagenomes.

In general, the abundance of different groups of viruses was more variable in the mid-harbour compared to the nearshore samples. Sample similarity based on community composition was assessed using UPGMA clustering of a Bray-Curtis dissimilarity matrix and reinforced this contrast of the nearshore and mid-harbour sites (Figure 3a). Though the nearshore and mid-harbour samples did not form distinct clusters overall, the nearshore samples clustered more closely together than the mid-harbour samples. The community composition at the two sites clustered together on July 30^th^ and September 10^th^, but resolved to distinct clusters on August 13^th^, August 27^th^ and September 24^th^.

**Figure 3:**
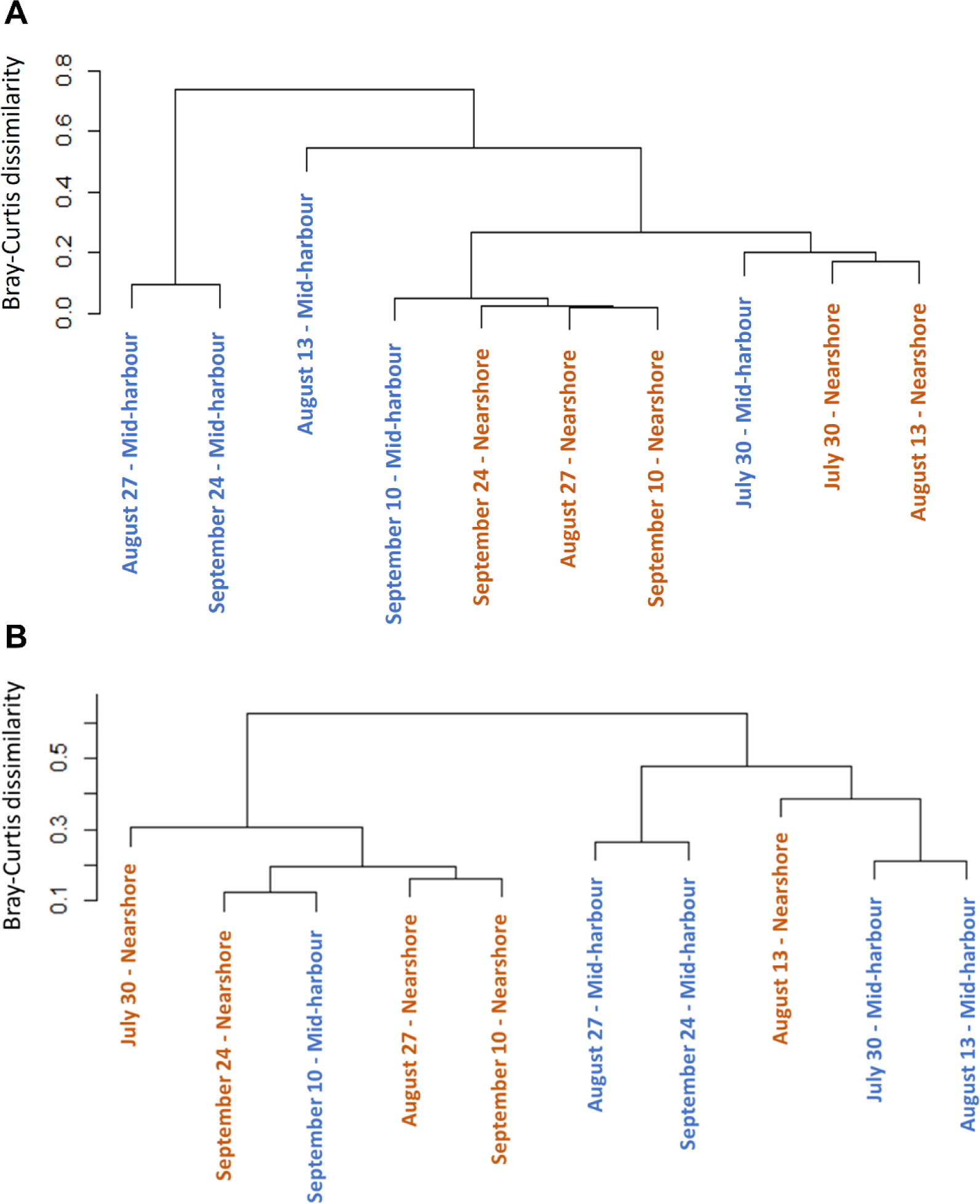
Dendrogram of cluster analysis based on Bray-Curtis dissimilarity index of the **a)** overall virus community and **b)** *Mimiviridae* community. Nearshore samples are coloured orange and mid-harbour samples are coloured blue.

### Mimiviridae community composition in Hamilton Harbour

Diverse *Mimiviridae* contigs were detected in Hamilton Harbour, including representatives of all subgroups and proposed subgroups. Most notably, the proposed ‘Klosneuvirinae’ subfamily [47] comprised a large proportion of the *Mimiviridae* community, ranging from 17.5% to 79.0% across all samples, and representing an average of 67.4% and 41.7% of the *Mimiviridae* community at the nearshore site and mid-harbour site, respectively. In general, the nearshore and mid-harbour sites appeared to host distinct *Mimiviridae* communities, a notion supported the Bray-Curtis dissimilarity clustering analysis (Figure 3b); samples from the two sites clustered separately, with the exceptions of September 10^th^ at the mid-harbour site and August 13^th^ at the nearshore site. In all nearshore samples and on September 10^th^ at the mid-harbour site, *Indivirus ILV1* were the most abundant representatives of the *Mimiviridae* community, while *Chrysochromulina ericinia* viruses (CeV) were the most abundant *Mimiviridae* in all mid-harbour samples except on September 10^th^ (Figure 4) when *Indivirus ILV1* were again dominant.

**Figure 4:**
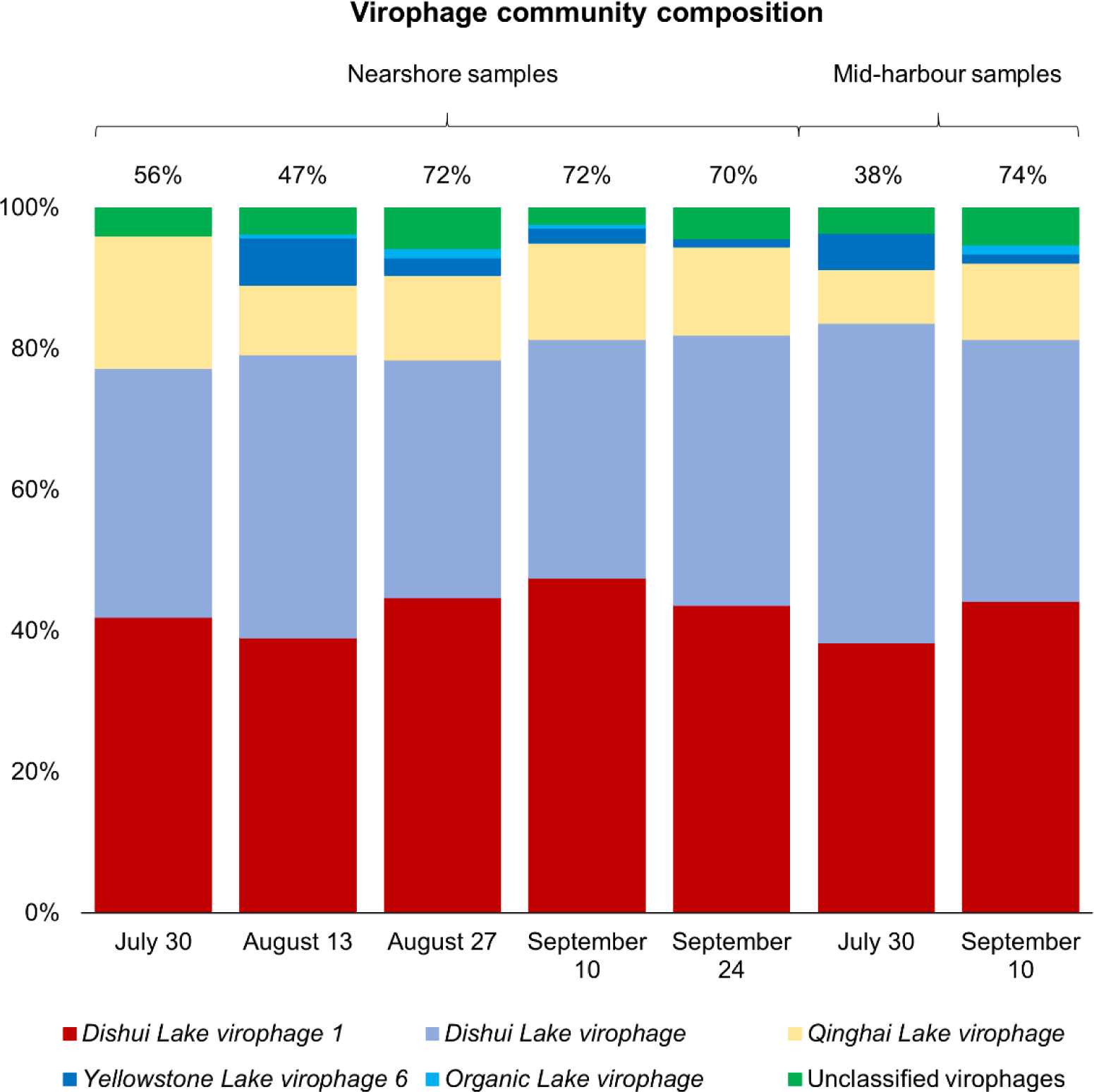
Virophage community composition in samples where virophages represent >10% of the total virus community. Numbers above bars are percentages of the total virus community that the entire bar represents.

Of the proposed subfamily ‘Mesomimivirinae’, Organic Lake phycodnavirus, Yellowstone Lake mimivirus and *Chrysochromulina ericinia* virus were the most abundant. The “Other *Mimiviridae*” category was created for *Mimiviridae* that were present in only a few of the samples and always at less than 2.0% of the community, and included the following Mesomimivirinae members: *Aureococcus anophagefferens, Phaeocystis pouchetii, Pyramimonas orientalis*, and *Phaeocystis globosa*. Also placed in the “Other *Mimiviridae*” category were *Acanthamoeba polyphaga* mimivirus, unclassified *Megaviridae*, Megavirus Iba, Powai Lake megavirus, and Mimivirus AB-566-O17, which were detected in 6 of the samples at less than 6.0% relative abundance.

### Virophage community composition in Hamilton Harbour

Given the similarity of virus community composition in the nearshore samples, we explored whether this consistency was upheld at the level of discrete taxa in the most abundant group, the virophages. Across all nearshore samples, virophage community composition was very similar regardless of date and contigs were annotated as five types of virophages: Dishui Lake virophage, Dishui Lake virophage 1, Yellowstone Lake virophage 6, Qinghai Lake virophage, and Organic Lake virophage (Figure 5). Across all samples, the majority of virophage contigs were most similar to Dishui Lake virophage 1 (43.4% average) and Dishui Lake virophage (36.3% average). Qinghai Lake virophages comprised 13.3% of the communities on average, and the Yellowstone Lake virophages were observed at lower abundances averaging only 2.5% of the virophage community. Least abundant were the Organic Lake virophages, which were detected at 0.5% abundance on average and were only detectable in 3 of the 5 nearshore samples. Unclassified virophage contigs were also detected at low abundances on all dates, averaging 4.0% of the virophage community. Overall, from July 30^th^ to September 24^th^ the virophage community composition at the nearshore site was remarkably similar. In contrast to virophage communities at nearshore sites, the mid-harbour samples from July 30^th^ and September 10^th^ were the only samples with virophage populations comprising >10% of the total virus community. Again, both samples were dominated by the Dishui Lake virophages (Dishui Lake virophage and Dishui Lake virophage), which comprised about 80% of the total virophage population. Regardless of site or date, the contigs most closely resembled virophages originally identified in Dishui Lake, China (Figure 5).

**Figure 5:**
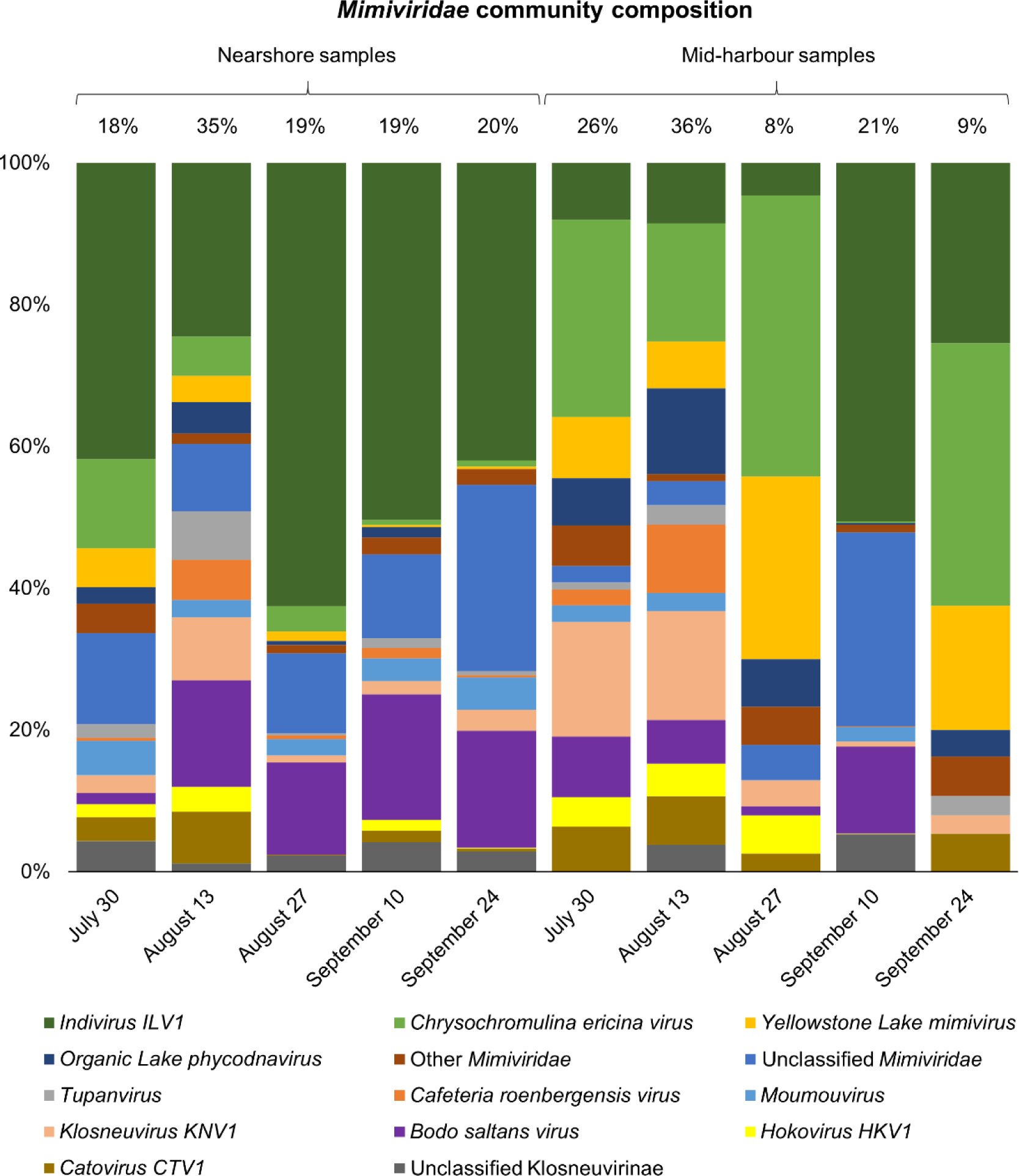
*Mimiviridae* community composition at the nearshore and mid-harbour sites from July 30^th^ to September 24^th^. Numbers above bars are percentages of the total virus community that the entire bar represents.

### Influence of environmental parameters on virus community composition

A canonical correspondence analysis (CCA) was performed to assess the influence of pH, temperature, redox potential, dissolved oxygen, Secchi depth and chlorophyll a on the viral groups at each site. The CCA model explained 47.8% (F = 7.12, Pr(>F) = 0.005) of the variability in the data. A test of 10,000 permutations revealed that chlorophyll a concentration was the only significant environmental parameter of those assessed. Temperature, pH, dissolved oxygen, Secchi depth, and redox potential were not significant explanatory factors of changes in virus community composition and relative abundances between samples. However, there was a strong inverse relationship of *Caudovirales* and chlorophyll a concentration. The August 27^th^ and September 24^th^ mid-harbour samples, which are dominated by *Caudovirales* (>70%), were similar in community composition (Figure 3a) and were negatively correlated with chlorophyll a. All nearshore samples clustered closely together and were positively correlated with chlorophyll a. Several parameters including ammonia, chloride, fluoride, sulfate, particulate organic carbon, particulate organic nitrogen, nitrate and nitrite, total dissolved nitrogen, total phosphorus, total dissolved phosphorus, total particulate phosphorus, and soluble reactive phosphorus were measured only on July 30^th^, September 10^th^ and September 24^th^ at both sites; however, none were significant explanatory variables of the virus communities on these dates. Interestingly, when the influence of environmental variables on the *Mimiviridae* community was assessed, chlorophyll a remained an explanatory variable, accounting for 37.8% of the variation in the *Mimiviridae* community (F = 4.85, Pr(>F) = 0.018). Of the parameters measured only on July 30^th^, September 10^th^ and September 24^th^, only particulate organic carbon (POC) significantly explained differences in the *Mimiviridae* communities between samples. In a separate CCA model, a test of 719 permutations revealed that POC accounted for 59.6% of the variation in the *Mimiviridae* community on July 30^th^, September 10^th^ and September 24^th^ (F = 5.90, Pr(>F) = 0.043).

## DISCUSSION

### Influence of databases

In this study we observed diverse virus communities that were both spatially and seasonally variable, and often dominated by virophages. However, as usual for environmental metagenomes, a large portion of sequences remained unclassified and were discarded at the annotation step of the data processing pipeline. This is especially the case for viruses, which have much fewer sequenced representatives in databases than prokaryotes or eukaryotes and are more likely to remain unannotated. For example, in May 2018, RefSeq release 88 contained >7500 viruses with >325,300 associated accessions (9648 nucleotide sequences and 315,742 protein sequences). In contrast, there were >50,400 bacteria with >100,583,400 associated accessions (12,367,951 nucleotide sequences and 88,193,695 protein sequences) (data from: ftp://ftp.ncbi.nlm.nih.gov/refseq/release/release-statistics/). An estimated <1% of the Earth’s virome has been discovered [48], and this underrepresentation in databases has a large influence on the reported diversity of viruses in metagenomes. The NCBI-nr database referenced in this study combines data from RefSeq as well as SwissProt, Protein Information Resource (PIR), Protein Databank (PDB), Protein Research Foundation (PRF), and GenBank. A combination of curated and non-curated sequences, the NCBI-nr database includes data from the assembly of environmental metagenomes including those deposited by private institutions. The use of this database as opposed to a curated database drastically increases the size of the reference database, increasing the likelihood of annotation, which is especially valuable for identifying taxa with few sequenced representatives.

Within viral sequence databases, certain groups of viruses have many sequenced representatives, while others are represented by only a few sequences. For example, there are >2000 complete *Caudovirales* genomes, but only 7 complete virophage (*Lavidaviridae*) genomes (data from https://www.ncbi.nlm.nih.gov/genomes/GenomesGroup.cgi; retrieved December 2018). This bias results in a higher chance of detecting *Caudovirales* than virophages, which are more likely to be unannotated and discarded in downstream analyses. Additionally, viral groups with more sequenced representation are more likely to have accurate annotations at the level of discrete taxa. Hence, the disproportionately high abundance of virophages detected in Hamilton Harbour samples compared to reference databases reinforces the discovery of their relatively high representation in this system, and likely other freshwater environments.

### Limitations of inferring virus presence and abundance from metagenomic datasets

This research infers the presence and abundance of viruses based on the relative abundances of assembled contigs from shotgun sequencing. It is important to note that the presence of contigs annotated as viruses may not necessarily indicate their presence. The intimate evolutionary relationship between viruses and their hosts has resulted in an abundance of viral genes present in host genomes. Unless viral homologues in host genomes are specifically annotated as such, genetic similarity searches may incorrectly annotate them as viruses [49]. However, upon examining the top 10 hits for individual contigs, there were few instances that contained a combination of bacteria and viruses; in most cases the top hits were all within the same virus family.

The concept of the “virocell” was introduced to distinguish between the living and non-living segments of the classical virus lifestyle [50]. The non-living portion of the virus lifecycle is the inactive virion that may become activated only if it contacts and infects a permissive host cell, while the living manifestation of a virus is the infectious and metabolically active intracellular portion of the virus lifecycle (i.e. the virocell). Counting viral contigs without distinguishing between active and inactive states may lead to an overestimation of the influence of viruses in a given environment if the data are not considered in light of this complex relationship between viruses and their hosts. Metagenomic research is limited by the inability to distinguish between viruses that are actively infecting hosts and virions that are inactive and simply adsorbed to cells or particles. Furthermore, in this study we could not distinguish between viruses integrated into host genomes and those present as virions. However, since DNA was extracted from the >0.22 um fraction rather than the traditional <0.22 um fraction, it is less likely that the virus contigs captured were derived from free virions in the water column. Most free virions, except some of the largest NCLDVs, would have passed through the filters, and only those that were adsorbed to cells or other particulate matter would have been captured during sample collection. Though virophages are small and could pass through the filters if present as free particles in the water column, it is also possible that several virophages could be adsorbed to individual giant virus hosts. For example, the virophage Sputnik is frequently observed to be situated within the fibrils of its *Mamavirus* host and is speculated to use these fibrils in order to gain entry into the viral host [51]. More research is required to confirm virophage host associations and methods of entry in order to adequately assess whether virophages commonly adsorb to their giant virus hosts, and what influence this may have on the interpretation of metagenomic data gathered in studies such as this. Regardless, the high abundance of virophages in the majority of Hamilton Harbour samples is indicative of their potential influence on the *Mimiviridae* and eukaryotic host communities in the harbour, and likely many other freshwater environments as well.

Though these observations contrast metagenomic studies in other freshwater lakes, it remains challenging to compare virus communities between studies due to differences in sampling, sequencing and data processing. Even the most widely-used analysis tools for metagenomic data yield different results [52], highlighting the need for standardized analysis tools and pipelines in order to facilitate ecologically relevant comparisons between studies.

### Physiochemical factors

Environmental variables that typically explain variation in virus communities on local scales, such as dissolved oxygen and temperature [6], were not significant explanatory variables of our data. This may be a reflection of virus dependence on host populations and the wide host ranges of virus families. For example, the *Mimiviridae* infect hosts ranging from heterotrophic protists like amoeba to photosynthetic protists, or algae, and because key resources for these eukaryotic microorganisms differ (DOC versus light), it might be anticipated that they respond differently to certain environmental factors. Nevertheless, the CCA model with chlorophyll a as a constrained variable explained 47.8% of the variation in the data, while other environmental parameters were insignificant. Therefore, it appears that fluctuations in algal biomass (inferred from chlorophyll a concentration) play a major role in shaping the virus communities in Hamilton Harbour. For example, chlorophyll a was inversely related to *Caudovirales* relative abundance, indicating that most *Caudovirales* contigs were derived primarily from phages of heterotrophic bacteria. The mid-harbour samples from August 27^th^ and September 24^th^ had undetectable or very low (0.1 µg/L) chlorophyll a concentrations, respectively, and the virus communities in these samples were dominated by *Caudovirales* and clustered closely together in both the CCA and the Bray-Curtis dissimilarity analyses. In contrast, the lowest chlorophyll a concentration noted for a nearshore sample was 2.7 µg/L, and the virus community in this sample (August 13^th^) was dominated by virophages and *Mimiviridae*. It is notable that the nearshore site is closer in proximity to wastewater effluents from Spencer Creek than the mid-harbour site. Although improvements have been made to the wastewater treatment plants surrounding Hamilton Harbour, effluents still contain phosphorus levels above targets set in the Hamilton Harbour Remedial Action Plan (HHRAP) [53]. These nutrient and organic carbon inputs might stimulate microbial growth at the nearshore site, in turn influencing virus communities.

The CCA of different *Mimiviridae* taxa against environmental parameters revealed that chlorophyll a remained a significant explanatory variable of the *Mimiviridae* community, reflecting their wide host range which includes both photosynthetic and non-photosynthetic hosts. For example, the algal virus CeV made up a large portion of the *Mimiviridae* community at the mid-harbour site, while *Indivirus ILV1*, a member of the proposed subfamily ‘Klosneuvirinae’ that were themselves assembled from metagenome sequences, was abundant in nearshore samples and is a suspected protist virus [47]. POC explained almost 60% of the variation in the *Mimiviridae* communities on July 30^th^, September 10^th^ and September 24^th^. Since POC data were only available for 6 of the 10 samples, it is unclear whether POC would remain a significant explanatory variable if the model included data from all samples.

Hamilton Harbour is known to be a highly variable system [54] that experiences regular seiches [55] and exchange flows with Lake Ontario, especially during the summer [27]. Circulation and mixing in the harbour are primarily controlled by prevailing winds [56]. Modelling of summer circulation patterns in Hamilton Harbour demonstrated the occurrence of a large eddy in the middle of the harbour at the location of our mid-harbour site, while the nearshore site was situated on the perimeter of a smaller eddy located at western end of the bay near Cootes Paradise [57]. The location of the sampling sites with respect to these eddies and Hamilton Harbour circulation may explain differences in virus community composition observed at the different sites on the same day. Hamilton Harbour is a major Canadian shipping port and large ships entering the harbour from Lake Ontario likely pass through the mid-harbour site on route to the port on the southern shore. Shipping traffic is one of the many factors potentially affecting virus community variability on the sampling dates that could not be considered in the present study. Given the high variability of Hamilton Harbour, it is perhaps unsurprising that the virus communities varied widely between sampling sites and dates. More unexpected was the relatively stable community throughout the mid-summer to late fall at the nearshore site, when bacterial and phytoplankton populations have been observed to be highly dynamic [58-60].

### The virophages

Virophages are small dsDNA viruses that co-infect eukaryotic hosts with giant dsDNA viruses [61]. Their co-infection has been shown to reduce giant virus fitness, thereby increasing survival of the cellular host [62,63]. Virophages were discovered only a decade ago [63], and little is known about their ecology. There is evidence to suggest that virophage-induced reduction of algal host mortality leads to longer and more frequent algal blooms [64], highlighting the ecological importance of virophages and their potential relevance to our eutrophic study site.

Virophage abundance in freshwater eutrophic lakes varies widely and has been reported as highly abundant in some environments [26], yet only detected at low abundances (<0.05%) [20], or even undetectable in other lakes [22]. Virophage abundances and distributions in lakes do not have clear seasonal or annual patterns of abundance, nor are there established relationships between dominant types of virophages in different environments [26]. In contrast to previous studies of viruses in eutrophic freshwater lakes, virophages were dominant in most samples from Hamilton Harbour. About 80% of the virophage community was annotated as Dishui Lake virophages in 7 of 10 samples regardless of date or site. Dishui Lake, China is a eutrophic freshwater lake that is more similar to Hamilton Harbour than the lakes where other types of virophages were observed such as Organic Lake, an Antarctic hypersaline lake, Qinghai Lake, a saline endorheic basin, or Yellowstone Lake, a lake that receives geothermal inputs. Due in part to their relatively recent discovery, sequence databases are limited with respect to the amount of information available for virophages, hence, specific virophage annotations in our samples may not be accurate. It is likely that the large portion of the virophage community annotated as Dishui Lake virophages are in fact a diverse array of virophages. The breakdown into discrete taxa did, however, reveal which virophages in reference databases were most similar to those detected in our dataset.

Other considerations that may impact the accuracy of virophage annotations include the presence of polintons/polintoviruses, polinton-like viruses and transpovirons (reviewed in [65]). Related to virophages, polintons/polintoviruses are self-synthesizing transposons that have genes homologous to virophages and giant viruses and are frequently integrated into eukaryotic genomes. Polinton-like viruses (PLV) are related to polintons and are commonly integrated into the genomes of green algae [65]. Mimiviruses can also be infected by transpovirons, which are small, dsDNA parasites that share homologous genes with virophages. During mimivirus reproduction, numerous transpovirons can accumulate within the host cytoplasm, within mimivirus particles, and within virophage particles. They also have the ability to integrate into the genomes of mimiviruses [66]. Considering their evolutionary relatedness and the presence of homologous genes in conjunction with the limitations of available databases, it would not be surprising to find that some of the virophages detected in our samples may in fact be polintons/polintoviruses, PLVs, or transpovirons. Further exploration and sequencing of virophage, polinton, PLV and transpoviron diversity is required to expand databases and improve accuracy of identification of these entities in metagenomic datasets. As a final comment on virophages, recent genome sequencing of a giant virus isolated from Lake Ontario revealed the presence of three putative virophages [67]. This giant virus and associated virophages infect the freshwater haptophyte *Chrysochromulina parva*; the *C. parva* virus is a close relative to the Hamilton Harbour mimivirus contigs annotated as *Chrysochromulina ericinia* virus. This observation supports the notion that the contigs assembled from Hamilton Harbour represent bona fide mimivirus-parasitizing virophages.

### Ecological relevance

The importance of viruses in aquatic environments is well documented. They are estimated to kill approximately 10% of the phytoplankton population and up to 50% of the bacterial population in surface marine waters, with greater impacts in high nutrient environments [68]. Especially in eutrophic aquatic systems, viruses are more active and are hypothesized to control host abundance, respiration and production [69]. They drive host community succession by targeting and lysing abundant members of the community, allowing less competitive species to thrive [70], and influencing fluctuations in dissolved and particulate organic matter pools. Viruses also act as key gene transfer agents that permit host adaptation and drive the evolution of microbial communities [71].

While marine viruses have been studied extensively over the past decade, freshwater viruses have received relatively little attention. Fundamental aspects of freshwater virus ecology, such as their distribution and patchiness in freshwater environments remain unknown. To our knowledge, this research is the first report of virus communities in Hamilton Harbour, and one of the few studies of virus communities in the Great Lakes. On some dates (e.g. September 24^th^) virus community composition was very different over the 3-km distance between the sites, while on other dates (e.g. September 10^th^), the community composition was very similar. At the level of virus families, relative abundances fluctuated at the mid-harbour site much more than the nearshore site, highlighting the high diversity of viruses on local scales and the impacts of small-scale environmental differences.

Since the recent discoveries of the *Mimiviridae* and their virophages, few studies have looked at the relative abundances of these groups over the duration of a season. We captured fluctuations in the *Mimiviridae* community over the summer and observed vastly different communities on the same date at different locations in the harbour. The stability of the virophage community abundance throughout the nearshore samples despite the changing *Mimiviridae* community was unexpected as *Mimiviridae* are the only known virus hosts of virophages [72]. While the nearshore and mid-harbour samples generally contained distinct *Mimiviridae* communities, the virophage community composition remained consistent. This highlights the complexity of these ecological relationships, and the gaps in our understanding of how these intimately associated viruses interact. The stability of virophage community composition despite the fluctuating Mimiviridae community appears to support the hypothesis that virophages and their *Mimiviridae* hosts are not connected solely by infection. The notion that virophages may enter eukaryotic host cells independently, remaining latent until infection by a giant virus, where they then compete with the giant virus for its replication machinery [72] is a possible explanation of the wide range of abundances in the virophage community while the *Mimiviridae* community remained relatively stable.

Given that Hamilton Harbour is known to support high algal biomass in the summer and early autumn months, it was surprising that the *Phycodnaviridae* were observed at low relative abundances in all samples collected for this study. Instead, *Mimiviridae* appeared to be the dominant algal viruses in Hamilton Harbour. The most common algae-infecting *Mimiviridae* were CeV, which were especially abundant at the mid-harbour site. The “Other dsDNA viruses” category did not contribute more than 4% in any individual sample and included mostly *Mimiviridae*. The small percentage of ssDNA viruses that were detected represent only those that were replicating in cells in the dsDNA form, since only dsDNA was targeted in the library preparation and sequencing. Therefore, the ssDNA viruses may be underrepresented compared to other virus groups which were detected as inactive virions adsorbed to particles in addition to active viruses replicating within cells.

## CONCLUSIONS

Overall, Hamilton Harbour metagenomes included a diverse array of viruses ranging from large dsDNA *Mimiviridae* to small ssDNA viruses. Relative abundances of virus families varied widely over relatively small spatial scales within the harbour, with higher consistency in the nearshore samples compared to the mid-harbour samples. Virophage relative abundances ranged widely across all samples and were the most abundant virus family in most samples. Though *Mimiviridae* are presumably intimately associated with virophages, their abundances did not fluctuate to the same extent as the virophages. A wide diversity of *Mimiviridae* were detected in the samples, and the two sites appeared to host distinct *Mimiviridae* communities. The abundances of discrete virophage taxa were remarkably stable despite the dissimilar *Mimiviridae* communities at the two sites, highlighting our limited understanding of how these intimately associated viruses interact. Equally unexpected was the low abundance of *Caudovirales* in most samples, contrasting other studies of freshwater virus communities. *Phycodnaviridae* abundances were also surprisingly low in all samples despite Hamilton Harbour’s capacity to support high algal biomass during the summer and autumn months, suggesting that *Mimiviridae* are the dominant algae-infecting viruses in this system. These findings provide insight into virus community structures in freshwater environments, expanding the documented diversity of freshwater virus communities, highlighting the potential ecological importance of virophages, and revealing distinct communities over small spatial scales.

## SUPPLEMENTARY MATERIALS

**Table S**.**1:** List of original taxonomic annotations and re-assigned taxonomic affiliations of virus contigs in Hamilton Harbour metagenomes.

## ACKNOWLEDGMENTS

This work was supported in part by an NSERC Discovery Grant (#RGPIN-2016-06022) awarded to S.M.S. and an Ontario Graduate Scholarship (OGS) awarded to C.N.P. Hamilton Harbour sampling, DNA extraction and metagenomic sequencing was funded by Environment Canada. Environmental parameters were collected and measured by researchers at York University (Dr. Lewis Molot’s group) and Environment Canada (Dr. Susan Watson’s group). These samples were processed by the staff at the National Laboratory for Environmental Testing (NLET). Our appreciation to Dr. Daniel Huson for correspondence regarding data processing in DIAMOND and MEGAN6-LR.

## AUTHOR CONTRIBUTIONS

Conceptualization, C.N.P. and S.M.S.; Methodology, C.N.P., R.R.F., R.S., and S.M.S; Validation, C.N.P. and S.M.S.; Formal Analysis, C.N.P. and S.M.S; Investigation, C.N.P., R.R.F., R.S., and S.M.S.; Resources, R.R.F. and S.M.S.; Data Curation, C.N.P. and S.M.S.; Writing – Original Draft Preparation, C.N.P. and S.M.S; Writing – Review & Editing, C.N.P., R.R.F., R.S., and S.M.S.; Visualization, C.N.P. and S.M.S.; Supervision, R.R.F. and S.M.S.; Project Administration, R.R.F. and S.M.S.; Funding Acquisition, R.R.F. and S.M.S.

## CONFLICTS OF INTEREST

The authors declare no conflicts of interest.

