## Supplemental Table 1 for "Metagenomic analysis of virus diversity and relative abundance in a eutrophic freshwater harbour"

SUPPLEMENTARY MATERIALS

**Table S.1:** List of originally assigned taxonomic annotation and re-assigned taxonomic grouping of virus contigs in Hamilton Harbour metagenomes

| **Original Annotation** | **Re-assigned Grouping** |
|---|---|
| root;**Viruses;** | Unclassified viruses |
| root;Viruses;**ssRNA viruses…** | Unclassified viruses |
| *Unclassified viruses* | |
| root;Viruses;unclassified viruses;**Pacmanvirus A23;** | Other dsDNA viruses |
| root;Viruses;unclassified viruses;**Faustovirus;** | Other dsDNA viruses |
| root;Viruses;unclassified viruses;**Sewage-associated circular DNA virus-1;** | Unclassified ssDNA viruses |
| root;Viruses;unclassified viruses;**Skeletonema virus LDF-2015a;** | Unclassified ssDNA viruses |
| root;Viruses;unclassified viruses;**Lake Sarah-associated circular virus-1;** | Unclassified ssDNA viruses |
| *dsDNA viruses, no RNA stage* | |
| root;Viruses;**dsDNA viruses, no RNA stage;** | Other dsDNA viruses |
| root;Viruses;dsDNA viruses, no RNA stage;**Iridoviridae;** | Iridoviridae |
| root;Viruses;dsDNA viruses, no RNA stage;**Poxviridae;** | Poxviridae |
| root;Viruses;dsDNA viruses, no RNA stage;**Marseilleviridae;** | Other dsDNA viruses |

| | |
|---|---|
| root;Viruses;dsDNA viruses, no RNA stage;**Tectiviridae;** | Other dsDNA viruses |
| *environmental samples <viruses>* | |
| root;Viruses;**environmental samples <viruses>;** | Unclassified viruses |
| root;Viruses;environmental samples <viruses>;**Organic Lake virophage;** | Virophages |
| root;Viruses;environmental samples <viruses>;uncultured environmental isolates;**uncultured Mediterranean phage uvDeep1…** | Unclassified bacteriophages |
| root;Viruses;environmental samples <viruses>;**uncultured marine virus;** | Unclassified viruses |
| root;Viruses;environmental samples <viruses>;**uncultured Mediterranean phage uvMED;** | Unclassified bacteriophages |
| root;Viruses;environmental samples <viruses>;**uncultured Mediterranean phage;** | Unclassified bacteriophages |
| root;Viruses;environmental samples <viruses>;**uncultured virus;** | Unclassified viruses |
| root;Viruses;environmental samples <viruses>;uncultured environmental isolates;**uncultured phage MedDCM-OCT-S04-C348;** | Unclassified bacteriophages |
| root;Viruses;unclassified bacterial viruses;environmental samples <bacteriophages>;**environmental Halophage eHP-25;** | Unclassified bacteriophages |
| *unclassified bacterial viruses* | |
| root;Viruses;**unclassified bacterial viruses;** | Unclassified bacteriophages |
| root;Viruses;unclassified bacterial viruses;**Agrobacterium phage Atu_ph07;** | Unclassified bacteriophages |
| root;Viruses;unclassified bacterial viruses;**Bradyrhizobium phage BDU-MI-1;** | Unclassified bacteriophages |

| | |
|---|---|
| root;Viruses;unclassified bacterial viruses;environmental samples <bacteriophages>;**Lake Baikal phage Baikal-20-5m-C28;** | Unclassified bacteriophages |
| root;Viruses;unclassified bacterial viruses;**Freshwater phage uvFW…** | Unclassified bacteriophages |
| root;Viruses;unclassified bacterial viruses;**Gordonia phage GMA2;** | Caudovirales - Siphoviridae |
| root;Viruses;unclassified bacterial viruses;**Methylophilaceae phage P19250A;** | Caudovirales - Siphoviridae |
| root;Viruses;unclassified bacterial viruses;**Ralstonia phage DU_RP_I;** | Caudovirales – Podoviridae? |
| root;Viruses;unclassified bacterial viruses;**Stenotrophomonas phage vB_SmaS-DLP_6;** | Caudovirales - Myoviridae |
| root;Viruses;unclassified bacterial viruses;**Synechococcus phage S-CAM7;** | Caudovirales - Myoviridae |
| root;Viruses;unclassified bacterial viruses;**Synechococcus phage S-CAM9;** | Caudovirales - Myoviridae |
| root;Viruses;unclassified bacterial viruses;**Lactobacillus phage Semele;** | Unclassified bacteriophages |
| root;Viruses;unclassified bacterial viruses;**Pontimonas phage phiPsal1;** | Unclassified bacteriophages |
| root;Viruses;unclassified bacterial viruses;**Pseudomonas phage pf16;** | Caudovirales - Myoviridae |
| root;Viruses;unclassified bacterial viruses;**Paracoccus phage Shpa;** | Caudovirales - Siphoviridae |
| root;Viruses;unclassified bacterial viruses;**Synechococcus phage S-EIVl;** | Caudovirales - Unclassified |
| root;Viruses;unclassified bacterial viruses;**Alteromonas phage PB15;** | Caudovirales - Siphoviridae |
| root;Viruses;unclassified bacterial viruses;**Acidovorax phage ACP17;** | Caudovirales - Myoviridae |
| root;Viruses;unclassified bacterial viruses;**Streptomyces phage BRock;** | Caudovirales - Myoviridae |
| root;Viruses;unclassified bacterial viruses;**Mycobacterium phage B1;** | Caudovirales |

| | |
|---|---|
| root;Viruses;unclassified bacterial viruses;**Erwinia phage vB_EamM_Caitlin;** | Caudovirales - Myoviridae |
| root;Viruses;unclassified bacterial viruses;**Vibrio phage 1.046.O._10N.286.52.E3;** | Unclassified bacteriophages |
| root;Viruses;unclassified bacterial viruses;**Xanthomonas phage XacN1;** | Caudovirales - Myoviridae |
| root;Viruses;unclassified bacterial viruses;**Nostoc phage A1;** | Caudovirales - Myoviridae |
| root;Viruses;unclassified bacterial viruses;**Pseudoalteromonas phage PHS21;** | Unclassified bacteriophages |
| *unclassified dsDNA phages* | |
| root;Viruses;dsDNA viruses, no RNA stage;**unclassified dsDNA phages;** | Unclassified bacteriophages |
| root;Viruses;dsDNA viruses, no RNA stage;unclassified dsDNA phages;**Idiomarinaceae phage 1N2-2;** | Unclassified bacteriophages |
| root;Viruses;dsDNA viruses, no RNA stage;unclassified dsDNA phages;**Methylophilales phage HIM624-A;** | Caudovirales – Podoviridae |
| root;Viruses;dsDNA viruses, no RNA stage;unclassified dsDNA phages;**Salicola phage CGphi29;** | Unclassified bacteriophages |
| root;Viruses;dsDNA viruses, no RNA stage;unclassified dsDNA phages;**Cyanophage KBS-S-2A;** | Caudovirales - Siphoviridae |
| root;Viruses;dsDNA viruses, no RNA stage;unclassified dsDNA phages;**Synechococcus phage S-CBP3;** | Caudovirales – Podoviridae |
| *unclassified dsDNA viruses* | |
| root;Viruses;dsDNA viruses, no RNA stage;**unclassified dsDNA viruses;** | Other dsDNA viruses |
| root;Viruses;dsDNA viruses, no RNA stage;unclassified dsDNA viruses;**Emiliania huxleyi virus PS401;** | Phycodnaviridae |

| | |
|---|---|
| root;Viruses;dsDNA viruses, no RNA stage;unclassified dsDNA viruses;**Phaeocystis globosa virus 14T;** | Mimiviridae |
| root;Viruses;dsDNA viruses, no RNA stage;unclassified dsDNA viruses;unclassified archaeal dsDNA viruses;Haloviruses;**Halovirus HVTV-1;** | Unclassified bacteriophages |
| *Phycodnaviridae* | |
| root;Viruses;dsDNA viruses, no RNA stage;Phycodnaviridae;environmental samples <Phycodnaviridae>;**Organic Lake phycodnavirus;** | Mimiviridae |
| root;Viruses;dsDNA viruses, no RNA stage;Phycodnaviridae;environmental samples <Phycodnaviridae>;**Organic Lake phycodnavirus 1;** | Mimiviridae |
| root;Viruses;dsDNA viruses, no RNA stage;Phycodnaviridae;environmental samples <Phycodnaviridae>;**Organic Lake phycodnavirus 2;** | Mimiviridae |
| root;Viruses;dsDNA viruses, no RNA stage;Phycodnaviridae;unclassified Phycodnaviridae;**Aureococcus anophagefferens virus;** | Mimiviridae |
| root;Viruses;dsDNA viruses, no RNA stage;Phycodnaviridae;unclassifiedPhycodnaviridae;**Chrysochromulina ericina virus;** | Mimiviridae |
| root;Viruses;dsDNA viruses, no RNA stage;Phycodnaviridae;unclassified Phycodnaviridae;**Phaeocystis pouchetii virus;** | Mimiviridae |
| root;Viruses;dsDNA viruses, no RNA stage;Phycodnaviridae;unclassified Phycodnaviridae;**Pyramimonas orientalis virus;** | Mimiviridae |
| root;Viruses;dsDNA viruses, no RNA stage;Phycodnaviridae;Prymnesiovirus;unclassified Prymnesiovirus;**Phaeocystis globosa virus [16T];** | Mimiviridae |
